# Near real-time monitoring of wading birds using uncrewed aircraft systems and computer vision

**DOI:** 10.1101/2024.05.14.594154

**Authors:** Ethan P. White, Lindsey Garner, Ben G. Weinstein, Henry Senyondo, Andrew Ortega, Ashley Steinkraus, Glenda M. Yenni, Peter Frederick, S. K. Morgan Ernest

## Abstract

Wildlife population monitoring over large geographic areas is increasingly feasible due to developments in aerial survey methods coupled with the use of computer vision models for identifying and classifying individual organisms. However, aerial surveys still occur infrequently, and there are often long delays between the acquisition of airborne imagery and its conversion into population monitoring data. Near real-time monitoring is increasingly important for active management decisions and ecological forecasting. Accomplishing this over large scales requires a combination of airborne imagery, computer vision models to process imagery into information on individual organisms, and automated workflows to ensure that imagery is quickly processed into data following acquisition. Here we present our end-to-end workflow for conducting near real-time monitoring of wading birds in the Everglades, Florida, USA. Imagery is acquired as frequently as weekly using uncrewed aircraft systems (aka drones), processed into orthomosaics (using Agisoft metashape), converted into individual level species data using a Retinanet-50 object detector, post-processed, archived, and presented on a web-based visualization platform (using Shiny). The main components of the workflow are automated using Snakemake. The underlying computer vision model provides accurate object detection, species classification, and both total and species-level counts for five out of six target species (White Ibis, Great Egret, Great Blue Heron, Wood Stork, and Roseate Spoonbill). The model performed poorly for Snowy Egrets due to the small number of labels and difficulty distinguishing them from White Ibis (the most abundant species). By automating the post-survey processing, data on the populations of these species is available in near real-time (< 1 week from the date of the survey) providing information at the time-scales needed for ecological forecasting and active management.

## Introduction

The need for wildlife monitoring over large spatial extents is increasingly common in the study and management of populations and ecosystems (Weinstein 2018). Large-scale surveys can be conducted using airborne monitoring from either crewed aircraft (airplanes, helicopters) or uncrewed aircraft systems (UAS, aka drones; Gonzalez et al. 2016). There is a growing need for this type of monitoring to occur frequently to support active management and ecological forecasting (Dietze et al. 2018, White et al. 2019).

Collecting large scale data at high frequencies is challenging because crewed flights are expensive, low altitude crewed flights can be dangerous, and processing the imagery from both crewed and UAS flights into data on wildlife populations can be difficult and time consuming (Gonzalez et al. 2016). The technology for both acquiring imagery and computational approaches for converting that imagery into counts has improved dramatically over the past decade. Uncrewed aircraft systems allow the collection of high resolution data over relatively large extents at significantly reduced costs (Gonzalez et al. 2016, Hodgson et al. 2018).

Computer vision allows this imagery to be automatically processed into data on the location and attributes of individual organisms, resulting in more accurate and less time consuming counts (Hodgson et al. 2018, Weinstein 2018, Kabra et al. 2022). Despite these advances, most aerial surveys still occur infrequently, and there are often long delays between the acquisition of airborne imagery and its conversion into population monitoring data. This is due, in part, to the continued use of hand annotations by many survey efforts, and a lack of automation for leveraging existing computer vision models to quickly produce large-scale predictions (Kabra et al. 2022). Accomplishing near real-time monitoring over large scales relies on a combination of computer vision models to process drone imagery into data on the location and species identity of individual organisms, and workflow automation to ensure that once imagery is acquired it is processed immediately (Marsh and Kurfess 2023).

Wading bird monitoring in the Everglades provides an ideal use case for near real-time monitoring using UASs. The Everglades is a unique ecosystem that has been severely impacted by over 100 years of human activity (Sklar et al. 2005) and is now the focus of one of the largest and most expensive ecosystem restoration efforts in history (Clarke and Dalrymple 2003, Sklar et al. 2005). Large wading birds (orders Pelecaniformes and Ciconiiformes) are a key component of the ecosystem. A number of restoration benchmarks are based on their nesting colonies including: the abundance of key species, where colonies are located, and the timing and success of reproductive events (Crozier and Gawlik 2003, Frederick et al. 2009). Studying wading birds and evaluating restoration benchmarks at the scale of the Everglades requires large-scale individual counts of species across 8500 km^2^. The current approach to these surveys involves flying small crewed aircraft across the region once a month during the breeding season with a single observer estimating counts of species in each colony (Bancroft et al. 1994, Frederick et al. 1996, 2003) and taking digital photographs from the window of the aircraft for later hand counting on a computer. Due to the extensive, and therefore time-consuming nature of hand counts, the only regularly reported data for this indicator is the annual maximum counts (where field observations are used to identify which month to count for each region of each colony for each species based on the peak breeding activity observed in the field).

While this annual data is crucial for understanding long-term trends in the recovery of the system, there is also a need for higher frequency data and models to inform active water management decisions and ecological forecasting in the ecosystem (Romañach et al. 2022).

Weekly data is ideal due to the speed at which the birds respond to changing environmental conditions and the needs of water managers (Romañach et al. 2022), but the cost and risk of crewed flights makes weekly surveys prohibitive. Even if more crewed flights were feasible, the time required to hand count photos means that most images would remain uncounted.

Hand-counted photos are also known to be error prone, with average error rates of up to 50% and a general tendency to underestimate the number of birds (Frederick et al. 1996, 2003); a problem that would likely become worse as the scale of hand counts increased. While using UAS imagery and computer vision tools eliminates the need for hand counting, weekly surveys of multiple locations still generates a nearly daily need to process imagery into data, which is costly and time consuming if a person is required to perform each of the multiple steps in that process.

Here, we describe the development of a large-scale monitoring system for the Everglades that combines UAS surveys for capturing images of entire colonies of wading birds, computer vision algorithms for detecting birds in imagery and identifying them to species, and automated workflows to ensure that imagery is quickly converted into actionable information on bird populations (Figure 1). This end-to-end system produces accurate counts for most species and has allowed us to conduct weekly monitoring of 8 colonies over the last 4 years, with less frequent data on an additional 32 colonies. Most of the system is fully automated, allowing images from the field to be converted to data on the web in a matter of days, thus facilitating near real-time monitoring of the associated populations at large scales.

**Figure 1.**
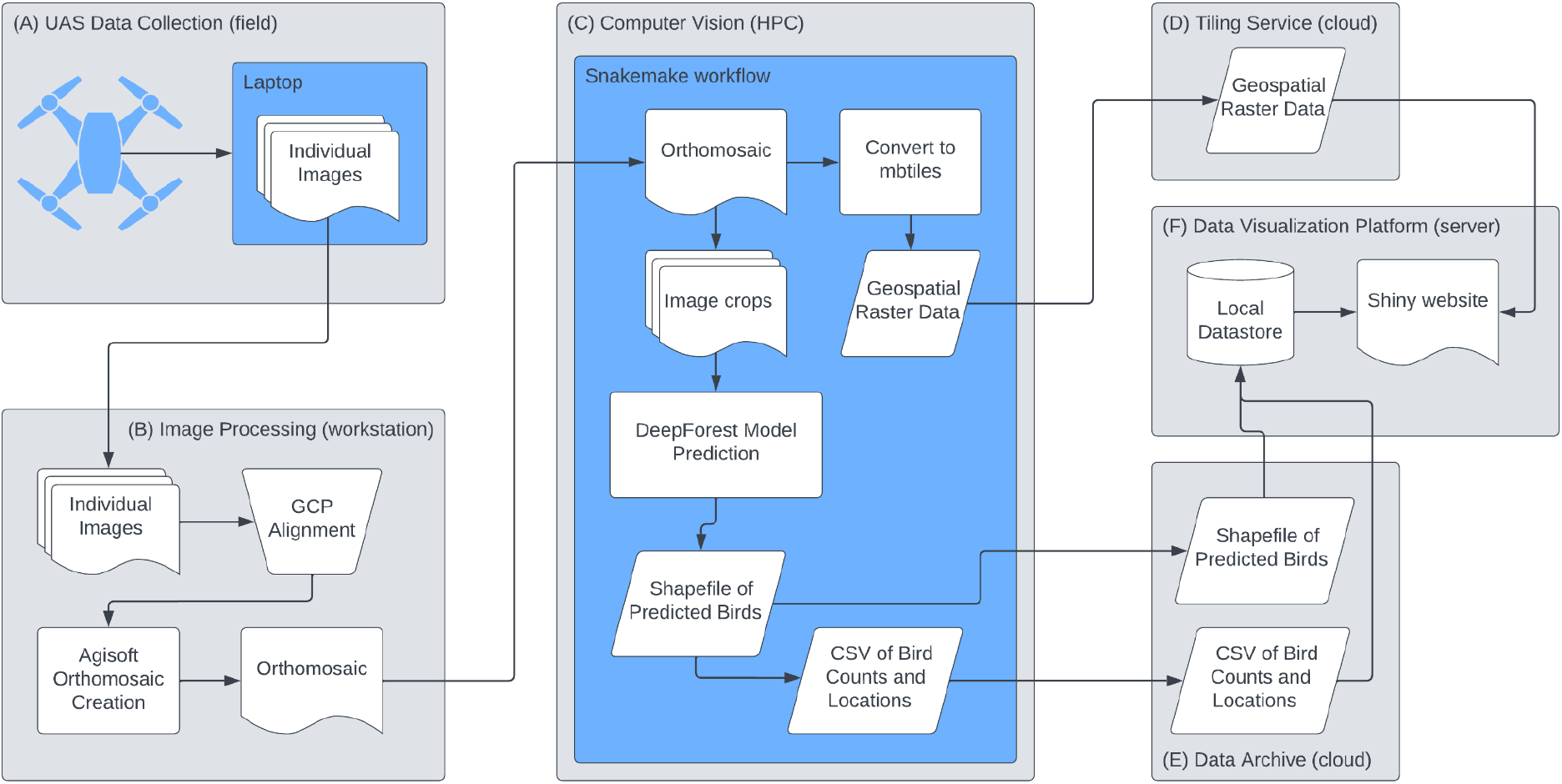
A diagram showing the full workflow for near real-time monitoring of Everglades wading birds involving five major components: (A) collection of raw imagery in the field using an uncrewed aircraft system; (B) processing imagery into orthomosaics on a workstation; (C) using an automated workflow to run a variety of computer vision and data processing steps on a high performance computing platform; (D) moving imagery to a cloud tiling service for data visualization; (E) archiving data on the location and counts of predicted birds; and (F) presenting these data and associated visualizations on the web.

## Materials and Methods

### Uncrewed aircraft system surveys

We have conducted ongoing UAS surveys over a total of 40 different active wading bird colonies in the Everglades starting in 2020. Surveys associated with this paper were conducted between February and May of 2020, 2021, 2022, and 2023. Breeding colonies in the Everglades vary considerably in both location and size across years. Therefore, in each year we selected a subset of colonies to survey more intensively with weekly surveys focused on large colonies that are consistently present across years and include a diversity of species (Figure 1A). We used the DJI Inspire II quadcopter, which is a robust off-the-shelf UAS platform designed for high-quality aerial photography and cinematography. The Inspire II was fitted with a DJI Zenmuse X7 24MP camera and a 35 mm equivalent lens with an integrated gimbal to provide image stabilization.

Repeatable 3D mapping flight plans were created in the DJI Ground Station Pro App mission planner. To obtain a ground sampling distance of approximately 1.0 cm, images captured via UAS were from an altitude of 76-92m AGL, shot at an angle of 15^°^ from nadir to obtain a slight profile of nesting birds. Images included >75% overlap in all four directions for orthomosaic production.

For the colonies surveyed weekly, we used ground control points (GCPs), large georeferenced targets easily visible in aerial imagery, to increase the accuracy of the real-world (latitude and longitude) positions in the orthomosaics. We placed 5 to 6 46cm × 46cm checkerboard patterned GCP targets, with easily identifiable center points, evenly around the colony perimeter. The precise latitude, longitude, and elevation of the center of each GCP was determined using a Trimble VRS network RTK solution coupled with the Florida Permanent Reference Network (a network of RTK reference stations; see Berber and Arslan 2013). In post-processing, the center of each target was identified in UAS imagery and the known geolocation used to georectify the resulting orthomosaic. This provided weekly real-world locations of birds within the colonies, which is necessary for tracking individual nests over time.

### Image processing

A UAS survey of a single colony generates tens to thousands of overlapping images stored in JPEG format. Using the AgiSoft Metashape Software, the images from each survey are combined into a single integrated image, known as an orthomosaic, to provide a complete view of the entire colony (Figure 1B). AgiSoft uses georeferencing information embedded in image metadata to align images. It then creates a digital elevation model using dense point cloud data and produces an orthorectified stitched image using this elevation model as its base map for depth rendering. We use camera accuracy settings of 10 meters for coordinate accuracy and 10 degrees for rotation accuracy. For sites with ground control points, the location data is loaded and the location markers are manually checked to ensure that they are placed in the center of the ground control point. Dense point clouds are created using medium quality (a 1:4 downscale factor) to optimize processing time and computer resources and aggressive depth filtering for outlier removal, the recommended setting for aerial data processing (Agisoft LLC 2024). This process produces a single GeoTIFF file for each survey of each colony. The resulting orthomosaic is manually checked to ensure that the individual photos are aligned well and modifications made to the default settings if necessary. The resulting files are large, so we use LZW compression with PREDICTOR=2 to produce lossless compression (reducing the space required to store the image without any loss of detail) that is efficient in the presence of spatial autocorrelation. This reduces the resulting file sizes by 2-3x, but many are still >4 GB so they are stored in BigTIFF format.

### Image annotation

Human labels for training the computer vision model were generated using a pragmatic combination of annotation approaches focusing on seven species: White Ibis (*Eudocimus albus*), Great Egret (*Ardea alba*), Great Blue Heron (*Ardea herodias*), Snowy Egret (*Egretta thula*), Wood Stork (*Mycteria americana*), Roseate Spoonbill (*Platalea ajaja*), and Anhinga (*Anhinga anhinga*). Our primary approach was to label birds in 3145 1500×1500 pixels crops from surveys conducted in 2020 and 2021. Crops were automatically extracted from the orthomosaics for each survey. For most crops every bird in each crop was annotated as a point location (the center of the bird) using one of the 7 species labels. Points were converted to bounding boxes by centering a box on each point using a 50×50 pixel box for larger species (Great Egret, Great Blue Heron, Wood Stork, Roseate Spoonbill, Anhinga) and a 36×36 pixel box for smaller species (White Ibis, Snowy Egret). We added a small number of additional crops for poorly represented classes by: 1) selecting imagery for annotation using a combination of expert field knowledge; and 2) running a preliminary model based on the primary crop data over survey imagery and identifying crops that were likely to include less common species. These crops were then annotated with bounding boxes and associated species labels. Images from the crop-based labels were split into separate datasets for training the model (∼90% of annotations) and testing its performance (∼10% of annotations).

Our secondary approach was to supplement the crop-based labels with labels generated from colony counts from 2022. Every bird in an opportunistically selected subset of colony surveys was annotated as a point location with a species label using Photoshop. Due to the scale of labeling, points were not placed carefully on the center of the bird, so to convert them into bounding boxes we used the bird detector module in the DeepForest Python package (Weinstein et al. 2022) to predict the location of birds within the image and associated the species label from each photoshop point with the overlapping bounding box. For the small number of points that did not have an overlapping bounding box we used a 1×1 m box centered on the point. These colony count labels were added to the training data, resulting in a total of 5128 image crops, including 50491 birds, for model training.

Test data for evaluating the performance of the model was based exclusively on the crop-based labels. To ensure that the test data was as accurate as possible, all labels in the test set were checked and adjusted if necessary by the lead of the labeling effort (Lindsey Garner). To allow for instances when it was not possible for a human to label a bird confidently to species, the labels for birds in the test set that could not be identified by the labeling lead were changed to indicate that the species was unknown. This produced a test set of 197 image crops, containing 4113 birds, with 94 birds labeled as unknown. All labeled data and associated imagery are openly available (Garner et al. 2024). See Garner et al. (2024) for additional details on annotation.

### Computer vision model

To detect and classify birds in the imagery, we used a RetinaNet-50 one-stage object detection model implemented using the DeepForest Python package (Figure 1C; Weinstein et al. 2020). RetinaNet-50 is a convolutional neural network with joint detection and classification heads (to allow the model to simultaneously identify and classify birds to species) that incorporates feature pyramids for capturing complex multi-scale structures in imagery and uses focal loss to address class imbalance between foreground and background classes (Lin et al. 2020). The model was pre-trained to detect birds, but not identify them to species (by detecting a single “bird” class), on a range of ecosystems and species (Weinstein et al. 2022). This allowed the model to learn general bird detection features. Starting with the backbone and regression head from this general model, we trained a seven-class species detector using the training labels described above. To avoid issues with detecting background objects as birds when predicting outside of these labeled datasets, we also included a selection of different background images that contained no birds in the training dataset. We trained the model for 62 epochs on NVIDIA A100 GPUs using stochastic gradient descent with momentum of 0.8. We explored different combinations of hyperparameters and selected a minimum bounding box confidence threshold of 0.3 to be considered a valid prediction, a non-max suppression threshold of 0.5, a batch size of 8, an initial learning rate of 0.001, and an IOU threshold of 0.4.

### Model evaluation

We evaluated the computer vision models using four metrics: 1) precision (the proportion of objects that the algorithm identified that match a human label); 2) recall (the proportion of human labels that are identified by the algorithm; 3) overall accuracy as measured by F1-scores; and 4) comparison of the counts generated by the algorithm to the counts from human labels using the root mean square error (RMSE) and the coefficient of determination about the 1:1 line (R^2^) on both untransformed and square root transformed data. Square root transformation was used to help emphasize the relationship in image crops with fewer than 100 individuals; the majority of the image crops. Evaluations were conducted for both the model’s ability to detect and count birds regardless of species (i.e., to identify birds and count the total number of birds in an image) and at the species level (i.e., to accurately identify birds to species). Birds that could not be conclusively identified by human labelers were excluded from species level analyses (n = 94 birds out of 4113 test labels), as were Anhinga (because there were very few Anhinga labels and they were only included as a nuisance class for model training). We focus on direct measures of model accuracy, not the relative fit compared to other modeling approaches, because in this applied context our concern is whether accurate counts can be accomplished, not whether one model produces slightly higher metrics of model fit.

### Near real-time prediction

To facilitate the rapid use of this UAS-based monitoring we developed an integrated workflow to quickly move from UAS surveys in the field, through image processing and computer vision algorithms, to data on bird abundances and locations (Figure 1). Imagery from the field is synched over the cloud to a workstation where orthomosaics are created (see *Image processing*). Orthomosaic creation is largely automated, but requires a researcher to check and update ground control point locations prior to image processing and check the resulting alignment. As soon as the orthomosaic is complete it automatically syncs to the University of Florida’s High Performance Computing Center. An automated daily workflow (Weinstein et al. 2024) built using the Snakemake workflow management system (Köster and Rahmann 2012, Mölder et al.2021) then runs all of the necessary tasks to convert the orthomosaic into data on birds. The workflow identifies the new orthomosaics, reprojects them, splits them into crops, and runs the computer vision algorithm to detect and classify all birds to species. It stores the resulting predictions as both geospatial data and tables of species counts, then processes the orthomosaic into a format appropriate for web-based visualization (mbtiles; García et al. 2012). The workflow pushes the processed image data to a hosted tiling service (Mapbox) and the bird predictions to a git repository (https://github.com/weecology/everwatch-predictions/). Finally this new data is used to update web-based visualization tools (a Shiny app written in R) to display the location and species of birds identified by the models on top of the imagery, along with counts of birds of different species (https://everwatch.weecology.org/).

## Results

The computer vision model accurately detected birds in imagery with a precision (proportion of predicted birds that are associated with human labels) of 0.84 and a recall (proportion of human labeled birds that are detected by the model) of 0.86. As a result, the model provides highly accurate counts of the number of birds in an image (Figure 2; R^2^ = 91.5%; R^2^ = 97.7%).

**Figure 2.**
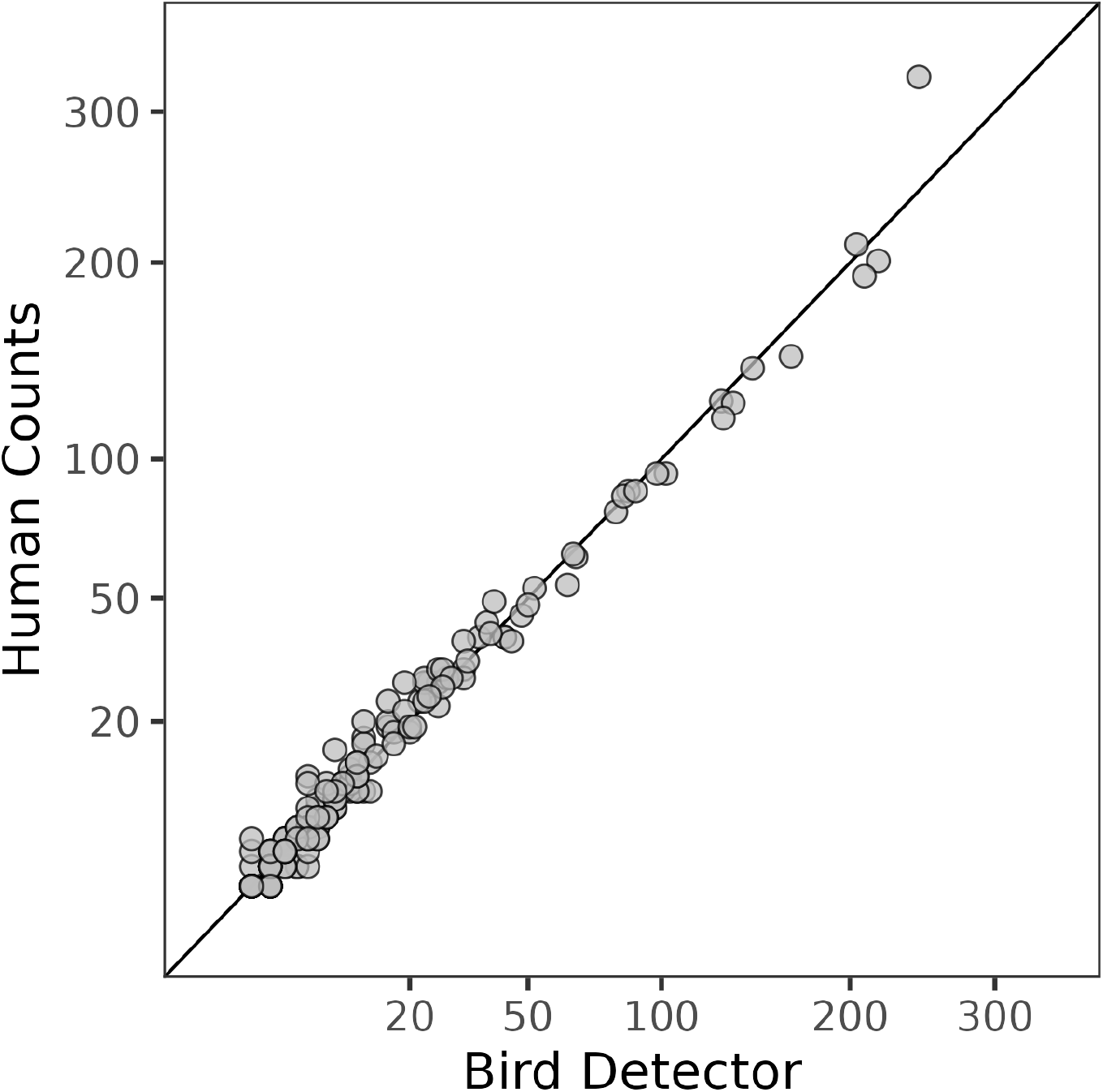
Comparison of the number of birds from human counts and the computer vision model for all species combined. The solid line is the 1:1 line indicating an exact match between human and computer vision counts. Axes are square root scaled to help show the relationship in image crops with fewer than 100 individuals (the majority of the image crops).

The machine learning model also performed well at classifying detected birds to the species level for most species (Table 1). Great Egrets, Roseate Spoonbills, White Ibis, and Woodstorks all have overall accuracies (F1-scores) of >0.85, precisions > 0.8, and recalls > 0.85. Great Blue Herons had lower, but still good values for F1 (0.75), precision (0.83), and recall (0.68). Great Blue Herons were confused with Great Egrets and Roseate Spoonbills (Figure 3). The model failed to correctly classify any Snowy Egrets resulting in no predictions for this species and zero or NA values for F1, precision, and recall (Table 1). Snowy Egrets are most commonly confused with White Ibis (Figure 3), a species that appears very similar when viewed in UAS imagery and is more common in the training data. Snowy Egrets are also confused with Great Egrets. Following standard computer vision convention, these species-level metrics are based on cases where the test bird was successfully detected by the model.

**Figure 3.**
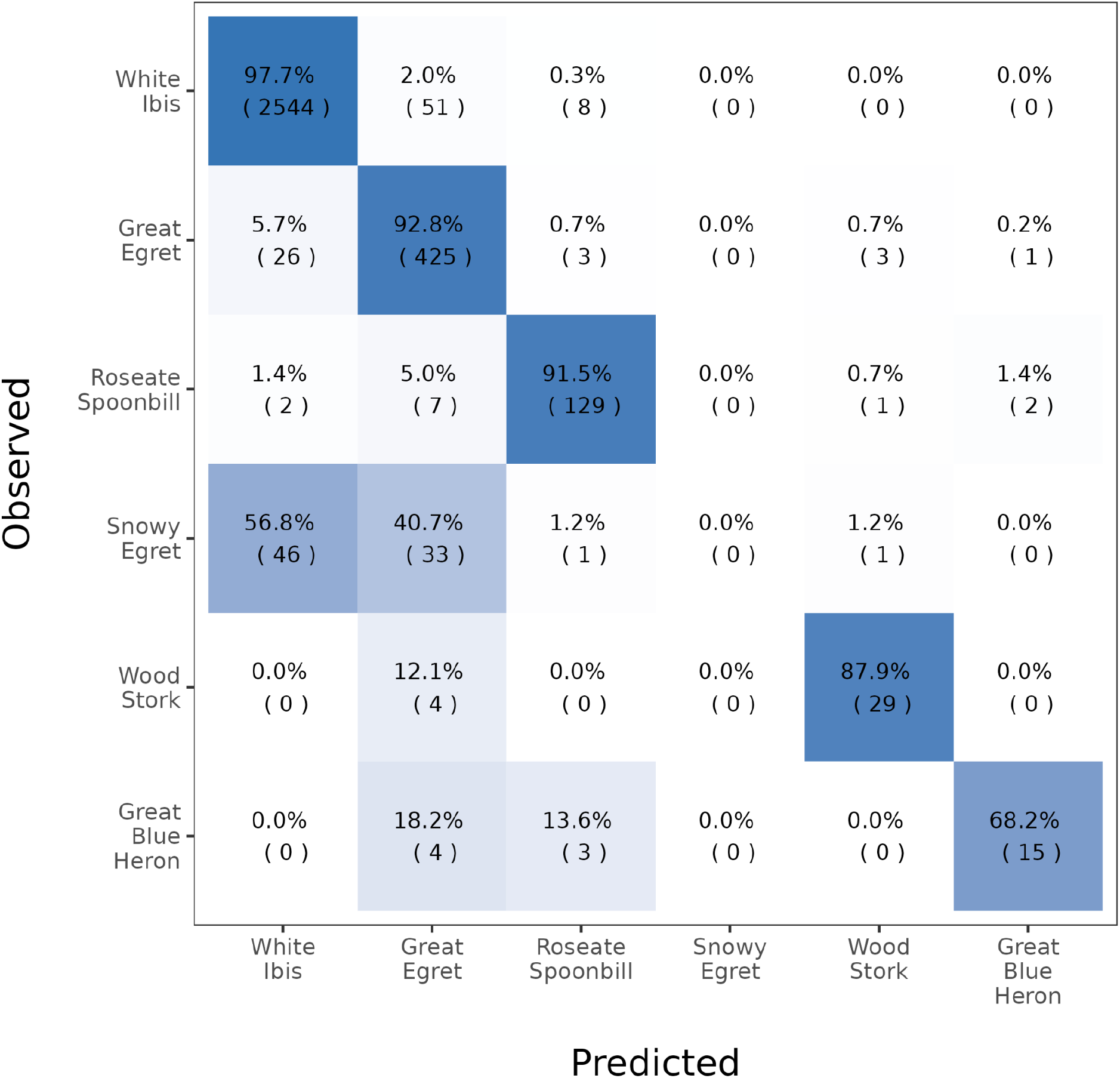
Confusion matrix showing correct and incorrect species classifications. All birds in each row belong to the species listed for that row and are predicted by the model to be the species in the associated column. Percentage values are the percentage of individuals for the observed species occurring in the cell (i.e, they are the counts in the cell divided by the sum of the counts over all columns in the same row times 100) and cells are colored by this percentage. Birds that were not detected by the model are not included. Species are ordered top to bottom and left to right based on their prevalence in the test data, with White Ibis being the most common and Great Blue Herons being the rarest.

**Table 1.** Accuracy (F1-score), precision, recall, coefficient of determination (R^2^) and root-mean-square error (RMSE) for the six focal species (Great Blue Heron, Great Egret, Roseate Spoonbill, Snowy Egret, White Ibis, and Wood Stork). Species are ordered top to bottom based on their prevalence in the test data, with White Ibis being the most common and Great Blue Herons being the rarest. Negative coefficients of determination indicate predictions that are worse than the mean abundance.

| **Species ($n_{\text{test}}$)** | **Accuracy (F1)** | **Precision** | **Recall** | **$R^2$** | **RMSE** |
| --- | --- | --- | --- | --- | --- |
| White Ibis (2603) | 0.97 | 0.97 | 0.98 | 0.98 | 5.95 |
| Great Egret (458) | 0.87 | 0.81 | 0.93 | 0.56 | 1.71 |
| Roseate Spoonbill (141) | 0.91 | 0.90 | 0.91 | 0.91 | 0.50 |
| Snowy Egret (81) | NA | NA | 0 | -0.11 | 2.45 |
| Wood Stork (33) | 0.87 | 0.85 | 0.88 | 0.94 | 0.37 |
| Great Blue Heron (22) | 0.75 | 0.83 | 0.68 | 0.35 | 0.38 |

The combination of high detection accuracy and good species classification results in accurate counts for most species (Figure 4). Root-mean-squared errors were low, with average errors of less than 2 individuals for all species except Snowy Egrets (for which the model broadly failed; see above) and White Ibis (Table 1). The White IBIS RMSE, 5.95, was expected to be higher due to the large number of White Ibis in each crop, and was also primarily driven by a single outlier. Coefficients of determination were also high, with the exception of Great Blue Herons and Snowy Egrets. Great Blue Herons are the least common species in the test set with only 0-3 individuals/crop. Counts this small make it difficult to obtain high values of R^2^ with discrete data.

**Figure 4.**
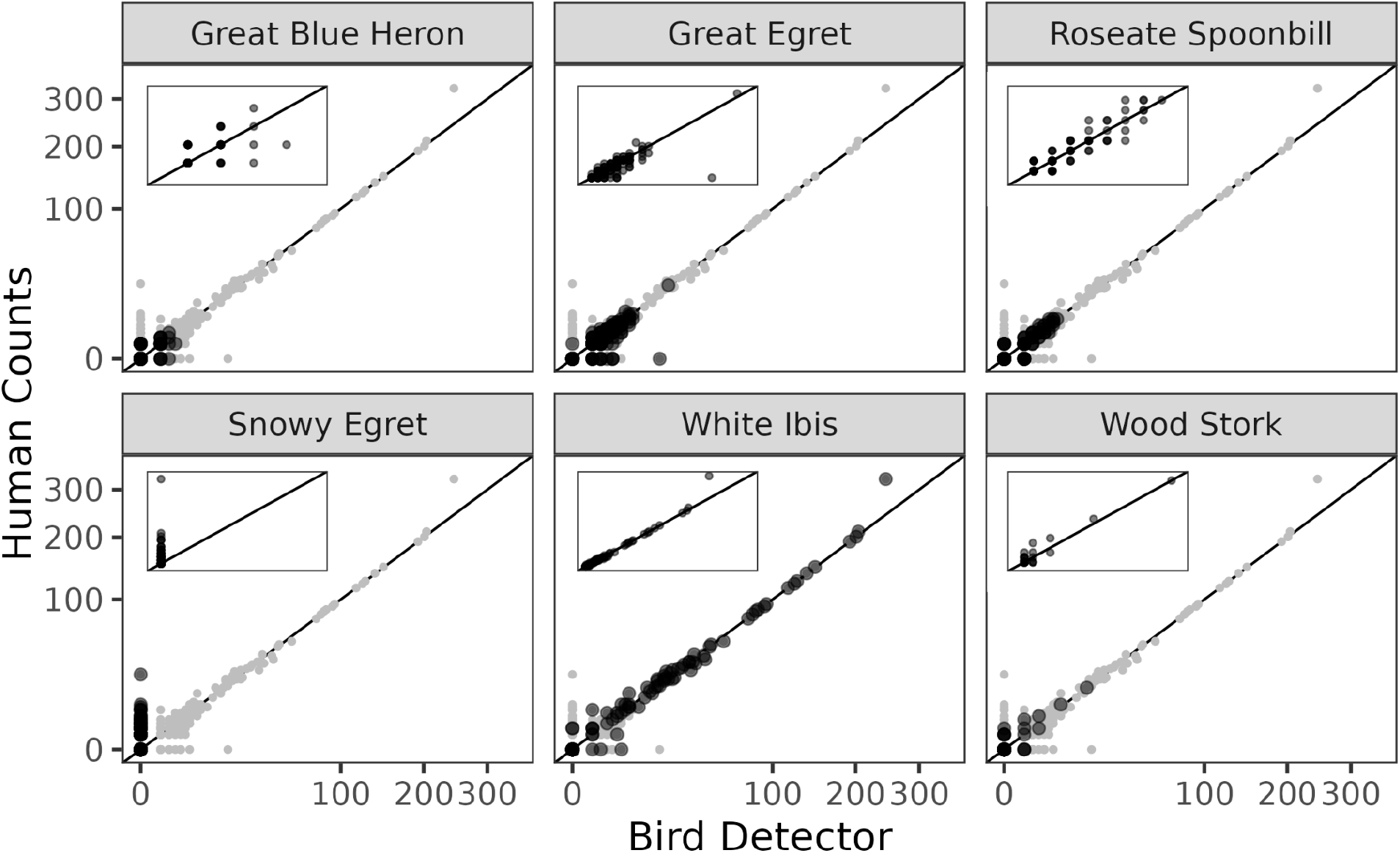
Comparison of counts of the six focal species (Great Blue Heron, Great Egret, Roseate Spoonbill, Snowy Egret, White Ibis, and Wood Stork) between human counts of birds in images and counts from the computer vision models. Dark gray dots are counts of the focal species in each crop. Light gray dots are counts of each of the other species in each crop. Solid lines are 1:1 lines indicating an exact match between human and computer vision counts. Axes in the main plots are square root scaled to help show the patterns in less common species. Insets show the relationship for only the focal species on unscaled (linear) axes.

The workflow system allows these predictions to be made in near real-time (∼2-7 days from the survey to available data). Results are displayed using web interfaces to allow researchers and managers to quickly explore the location, counts, and species of birds in the intensively surveyed colonies (Figure 5). While data are generated in near real-time, they are embargoed from the public until the end of the breeding season to limit their use in ecotourism in ways that could increase disturbance of the nesting colonies.

**Figure 5.**
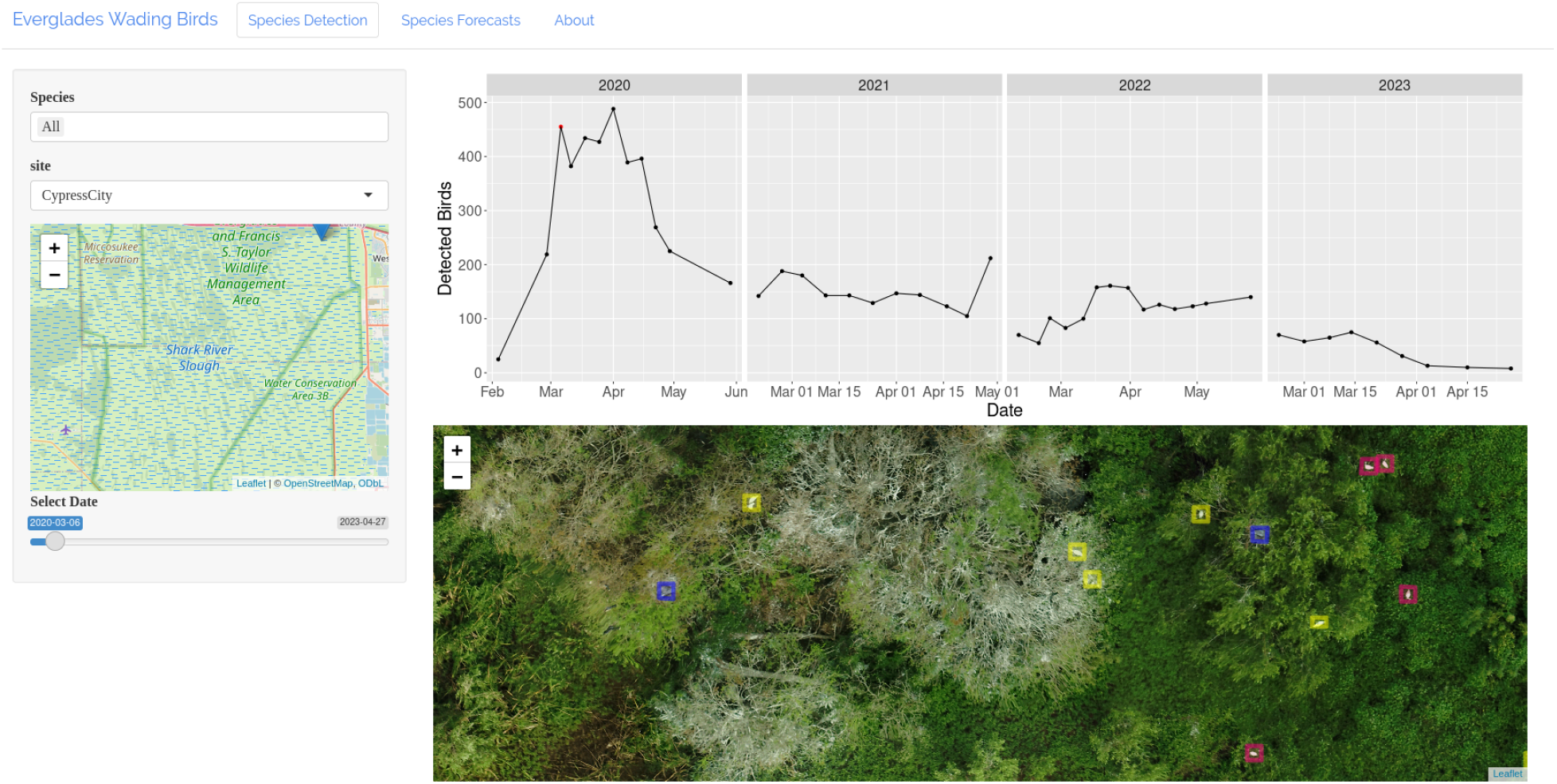
Screenshot of the https://everwatch.weecology.org/ website displaying imagery and predictions for birds in the Cypress City colony on March 6th, 2020. Three species are visible Roseate Spoonbills (pink boxes), Great Egrets (yellow boxes), and Great Blue Herons (blue boxes). Time-series of the total bird counts for each survey at the site are shown for the 2020-2023 breeding seasons.

## Discussion

We developed an end-to-end system using UAS surveys, computer vision, and workflow automation to facilitate near real-time monitoring of wading birds in the Everglades. Conducting this type of near real-time monitoring at large scales relies on computer vision models to transform UAS imagery into useful information on the location and species identity of birds, paired with workflow automation to ensure that once imagery is acquired it is quickly processed into data. The model we developed for Everglades wading birds performs well based on both common computer vision measures and the ecologically relevant measure of counts of individuals by species. The workflow system automates all but one step following the UAS survey, allowing rapid creation of data from imagery. This results in the availability of data on nesting populations within a week of the associated survey instead of waiting months for human labeling to be completed. This type of automated workflow could be broadly beneficial to the growing number of systems where large scale wildlife monitoring is needed and computer vision models have been shown to be effective (Lyons et al. 2019, Kellenberger et al. 2021, Kabra et al. 2022, Gayá-Vilar et al. 2024). These approaches will be particularly helpful in systems requiring higher frequency or larger scale monitoring (most wildlife applications still involve single flights over one or two locations; Hollings et al. 2018) and actively managed systems that require low latency access to monitoring data to support rapid decision making.

While this sort of approach can dramatically improve the speed and efficiency of monitoring efforts (Marsh and Kurfess 2023), computer vision models are imperfect and do not integrate external information in the same way as human annotators. Many of the mistakes the model makes are species misclassifications for images that are difficult even for humans because the birds are partially hidden by vegetation or distorted by the orthomosaic processing. Humans typically label these birds using additional context like the species identity of other birds in the surrounding area and knowledge from the field of what species were dominant at the location in the survey. While this sort of human inference is expected to be useful, there is no ground truth for these cases so the consequences of this type of human inference on accuracy are currently unknown. The current computer vision model cannot take into account these types of external information, but we are pursuing approaches to integrate spatial structure and expert knowledge in the future. To address the imagery distortions that are a natural part of the orthomosaic processing, we are exploring approaches to performing species classification in raw, undistorted imagery and then incorporating that information into an orthomosaic. Orthomosaic distortions are most common in imagery with large variation in vegetation height, which differentially affects species such as Wood Storks and Great Blue Herons that nest in tall trees.

Despite these limitations, our results show that for small image crops the model can achieve approximately human level accuracy, with very similar counts and instances of model confusion typically associated with birds that are difficult for humans to identify without additional context. This equivalent-to-human performance occurs in the context most advantageous to human annotators, where only a subset of a year’s worth of imagery is provided in small crops with small numbers of birds to count. The comparison to human labeling, therefore, provides a strong test of algorithm performance against the best possible conditions for human annotation efforts. In real-world contexts, hand counting imagery from every survey at the scale of our Everglades monitoring effort would be expensive and time consuming. We currently monitor half a dozen colonies approximately once a week, with the goal of being able to provide near real-time feedback on bird dynamics and reproduction to water managers. In addition to the aforementioned cost and time constraints, humans fatigue quickly at this type of task and therefore we expect human performance to degrade substantially at large scales resulting in high error rates (Frederick et al. 1996, 2003). Therefore, in the absence of large numbers of human annotators, the computer vision based counts are likely to be more accurate than human based efforts for identifying all of the birds in the weekly imagery due to the lack of the fatigue factor. This is supported by other imagery counting assessments that show that variability between human annotators is high in these contexts, meaning that at least some annotators are making significant mistakes (Bowler et al. 2020, Hayes et al. 2021).

The ability of computer vision models to efficiently count and identify wildlife at large scales does not replace the importance of field biologists with domain expertise. Instead, it reduces the amount of highly repetitive work that is necessary for monitoring, allowing field biologists to ask broader questions and focus their annotation work on checking and improving algorithmic approaches to counting. An ideal system for this type of monitoring would combine human expertise with the ability of computer vision models to consistently identify most individuals for an unlimited number of birds. Therefore, an important next step will be to build systems that start by labeling all birds based on computer vision and then involve human experts in: 1) reviewing those labels, aided by highlighting the birds that are mostly likely to be miscategorized by the model); 2) updating those labels based on expert knowledge and field notes (e.g., if the researcher knows from field observations that there was a flock of Snowy Egrets that was misclassified by the model as White Ibis); and 3) using active learning to select images for annotation that will lead to the best improvements in the future performance of the model.

While this approach has made it feasible to monitor a subset of colonies more frequently and efficiently than possible using conventional approaches, the current limitations of UAS make them insufficient for monitoring the entire Everglades. This is due to difficulties in reaching remote locations by airboat, regulations preventing UAS use in some areas (i.e., Everglades National Park and Loxahatchee National Wildlife Refuge), and the still time-consuming nature of large-scale UAS surveys. Many colonies still need to be surveyed using the conventional approach of human counts from small aircraft combined with pictures taken from the window of the plane that are manually aligned and counted by researchers. Fully replacing these laborious and time consuming manual counts requires the ability to acquire UAS-quality imagery at colonies across the entire study region. Airplanes equipped with cameras, which have a long history in wildlife monitoring (e.g., Conn et al. 2021), can collect imagery across large areas that could then be fed into automated workflows using models similar to ours but fine-tuned for plane-based imagery. On-board processing of imagery during flights may also be possible to filter and keep only images likely to contain target species, which would reduce the vast amounts of imagery collected during broad surveys. Satellite imagery is yet another platform for obtaining large-scale imagery of birds, especially when they occur in large clusters or colonies (Lynch et al. 2012, Fretwell et al. 2017). New model development would be required for detecting birds in satellite imagery, and increases in satellite resolution may be required before identification to species would be possible. Because high-resolution satellite imagery and plane flights are much more expensive than UAS surveys, we envision a future that combines low frequency, large-scale monitoring by satellites and/or planes for coarser resolution monitoring at infrequent intervals and UAS surveys for finer-scale information across locations of particular interest.

The tools for large scale monitoring in wildlife ecology, conservation, and management have advanced significantly through the use of UAS imagery and computer vision (Gonzalez et al. 2016, Hodgson et al. 2018, Weinstein 2018, Corcoran et al. 2021, Kabra et al. 2022). The majority of research in this area focuses on a single component of this task, typically either the UAS surveys or the performance of the computer vision models. While assessing each component of the process is valuable, fully leveraging these advances in technology to conduct near real-time monitoring of wildlife populations requires integrating individual components to produce end-to-end systems (Marsh and Kurfess 2023). These end-to-end systems allow imagery from the field to be quickly and efficiently converted into data that can be used to improve conservation and management decisions. We share our current system for performing wildlife monitoring as part of a major ongoing restoration effort and our vision of further improvements, with the hope of helping to encourage and guide the development and deployment of this type of system more broadly.

## Acknowledgements

This research was supported by grants from the Army Corps of Engineers (W912HZ-20-2-0022/3 to S.K.M. Ernest and P. Frederick), the South Florida Water Management District (4500126520 to S.K.M. Ernest and P. Frederick), the Gordon and Betty Moore Foundation’s Data-Driven Discovery Initiative (GBMF4563 to E.P. White), the National Science Foundation (DEB-2326954 to E.P. White and S.K.M. Ernest), and the USDA National Institute of Food and Agriculture via two Hatch Projects (FLA-WEC-005983 to S.K.M. Ernest and FLA-WEC-005944 to E.P. White). We thank Jerry Lorenz for critical logistical assistance that helped maintain our UAS program and Mark Cook for support and feedback on using UAS to monitor wading bird colonies. Finally we thank John Rouse and Cassandra Farley for helping us navigate the complex regulatory landscape surrounding the use of UAS in Florida.

## Author Contributions Statement

EPW, LG, BGW, AO, HS, GMY, PF, and SKME conceived the ideas and designed methodology; LG, AO, and AS collected the data; EPW, BGW, HS, and AO analysed the data; EPW and SKME led the writing of the manuscript. All authors contributed critically to the drafts and gave final approval for publication.

## Data Accessibility

All data and code associated with this paper are publicly archived on Zenodo including: 1) data for training the computer vision model (https://zenodo.org/doi/10.5281/zenodo.11165945); 2) code for training the computer vision model and producing the figures in this paper (https://zenodo.org/doi/10.5281/zenodo.11187070); 3) workflow code (https://zenodo.org/doi/10.5281/zenodo.11127010); 4) predictions produced for the location of birds (https://zenodo.org/doi/10.5281/zenodo.11191295); 5) Shiny app for data visualization (https://zenodo.org/doi/10.5281/zenodo.7004056).

## References

Agisoft LLC. 2024. Agisoft Metashape User Manual - Standard Edition, Version 2.1.

Bancroft, G. T., A. M. Strong, R. J. Sawicki, W. Hoffman, and S. D. Jewell. 1994. Relationships among wading bird foraging patterns, colony locations, and hydrology in the Everglades. Pages 615–658 The Everglades: The Ecosystem and Its Restoration. St. Lucie Press, Delray Beach, Florida, USA.

Berber, M., and N. Arslan. 2013. Network RTK: A case study in Florida. Measurement 46:2798–2806.

Bowler, E., P. T. Fretwell, G. French, and M. Mackiewicz. 2020. Using Deep Learning to Count Albatrosses from Space: Assessing Results in Light of Ground Truth Uncertainty. Remote Sensing 12:2026.

Clarke, A. L., and G. H. Dalrymple. 2003. $7.8 Billion for Everglades Restoration: Why Do Environmentalists Look So Worried? Population and Environment 24:541–569.

Conn, P. B., V. I. Chernook, E. E. Moreland, I. S. Trukhanova, E. V. Regehr, A. N. Vasiliev, R. R. Wilson, S. E. Belikov, and P. L. Boveng. 2021. Aerial survey estimates of polar bears and their tracks in the Chukchi Sea. PLOS ONE 16:e0251130.

Corcoran, E., M. Winsen, A. Sudholz, and G. Hamilton. 2021. Automated detection of wildlife using drones: Synthesis, opportunities and constraints. Methods in Ecology and Evolution 12:1103–1114.

Crozier, G. E., and D. E. Gawlik. 2003. Wading Bird Nesting Effort as an Index to Wetland Ecosystem Integrity. Waterbirds 26:303–324.

Dietze, M. C., A. Fox, L. M. Beck-Johnson, J. L. Betancourt, M. B. Hooten, C. S. Jarnevich, T. H. Keitt, M. A. Kenney, C. M. Laney, L. G. Larsen, H. W. Loescher, C. K. Lunch, B. C. Pijanowski, J. T. Randerson, E. K. Read, A. T. Tredennick, R. Vargas, K. C. Weathers, and E. P. White. 2018. Iterative near-term ecological forecasting: Needs, opportunities, and challenges. Proceedings of the National Academy of Sciences 115:1424–1432.

Frederick, P. C., B. Hylton, J. A. Heath, and M. Ruane. 2003. Accuracy and variation in estimates of large numbers of birds by individual observers using an aerial survey simulator. Journal of Field Ornithology 74:281–287.

Frederick, P. C., T. Towles, R. J. Sawicki, and G. T. Bancroft. 1996. Comparison of Aerial and Ground Techniques for Discovery and Census of Wading Bird (Ciconiiformes) Nesting Colonies. The Condor 98:837–841.

Frederick, P., D. E. Gawlik, J. C. Ogden, M. I. Cook, and M. Lusk. 2009. The White Ibis and Wood Stork as indicators for restoration of the everglades ecosystem. Ecological Indicators 9:S83–S95.

Fretwell, P. T., P. Scofield, and R. A. Phillips. 2017. Using super-high resolution satellite imagery to census threatened albatrosses. Ibis 159:481–490.

García, R., J. P. de Castro, E. Verdú, M. J. Verdú, L. M. Regueras, R. García, J. P. de Castro, E. Verdú, M. J. Verdú, and L. M. Regueras. 2012. Web Map Tile Services for Spatial Data Infrastructures: Management and Optimization. Page Cartography - A Tool for Spatial Analysis. IntechOpen.

Garner, L., B. Weinstein, M. Rickershauser, M. Baldino, H. Coates, M. Commins, T. Faber, J. Gula, S. Van Ert, E. White, and S. K. M. Ernest. 2024, May 13. EverWatch benchmark: training and evalutation data for detection and species classification of Everglades wading birds from airborne imagery. Zenodo.

Gayá-Vilar, A., A. Cobo, A. Abad-Uribarren, A. Rodríguez, S. Sierra, S. Clemente, and E. Prado. 2024. High-resolution density assessment assisted by deep learning of Dendrophyllia cornigera (Lamarck, 1816) and Phakellia ventilabrum (Linnaeus, 1767) in rocky circalittoral shelf of Bay of Biscay. PeerJ 12:e17080.

Gonzalez, L. F., G. A. Montes, E. Puig, S. Johnson, K. Mengersen, and K. J. Gaston. 2016. Unmanned Aerial Vehicles (UAVs) and Artificial Intelligence Revolutionizing Wildlife Monitoring and Conservation. Sensors 16:97.

Hayes, M. C., P. C. Gray, G. Harris, W. C. Sedgwick, V. D. Crawford, N. Chazal, S. Crofts, and D. W. Johnston. 2021. Drones and deep learning produce accurate and efficient monitoring of large-scale seabird colonies. Ornithological Applications 123:duab022.

Hodgson, J. C., R. Mott, S. M. Baylis, T. T. Pham, S. Wotherspoon, A. D. Kilpatrick, R. Raja Segaran, I. Reid, A. Terauds, and L. P. Koh. 2018. Drones count wildlife more accurately and precisely than humans. Methods in Ecology and Evolution 9:1160–1167.

Kabra, K., A. Xiong, W. Li, M. Luo, W. Lu, T. Yu, J. Yu, D. Singh, R. Garcia, M. Tang, H. Arnold, A. Vallery, R. Gibbons, and A. Barman. 2022. Deep object detection for waterbird monitoring using aerial imagery. Pages 455–460 2022 21st IEEE International Conference on Machine Learning and Applications (ICMLA).

Kellenberger, B., T. Veen, E. Folmer, and D. Tuia. 2021. 21 000 birds in 4.5 h: efficient large-scale seabird detection with machine learning. Remote Sensing in Ecology and Conservation 7:445–460.

Köster, J., and S. Rahmann. 2012. Snakemake—a scalable bioinformatics workflow engine. Bioinformatics 28:2520–2522.

Lin, T.-Y., P. Goyal, R. Girshick, K. He, and P. Dollár. 2020. Focal Loss for Dense Object Detection. IEEE Transactions on Pattern Analysis and Machine Intelligence 42:318–327.

Lynch, H. J., R. White, A. D. Black, and R. Naveen. 2012. Detection, differentiation, and abundance estimation of penguin species by high-resolution satellite imagery. Polar Biology 35:963–968.

Lyons, M. B., K. J. Brandis, N. J. Murray, J. H. Wilshire, J. A. McCann, R. T. Kingsford, and C. T. Callaghan. 2019. Monitoring large and complex wildlife aggregations with drones. Methods in Ecology and Evolution 10:1024–1035.

Marsh, P. K., and F. J. Kurfess. 2023. A software pipeline for automated wildlife population sampling. Frontiers in Conservation Science 4.

Mölder, F., K. P. Jablonski, B. Letcher, M. B. Hall, C. H. Tomkins-Tinch, V. Sochat, J. Forster, S. Lee, S. O. Twardziok, A. Kanitz, A. Wilm, M. Holtgrewe, S. Rahmann, S. Nahnsen, and J. Köster. 2021, April 19. Sustainable data analysis with Snakemake. F1000Research.

Romañach, S. S., S. M. Haider, C. Hackett, M. McKelvy, and L. G. Pearlstine. 2022. Managing multiple species with conflicting needs in the Greater Everglades. Ecological Indicators 136:108669.

Sklar, F. H., M. J. Chimney, S. Newman, P. McCormick, D. Gawlik, S. Miao, C. McVoy, W. Said, J. Newman, C. Coronado, G. Crozier, M. Korvela, and K. Rutchey. 2005. The ecological–societal underpinnings of Everglades restoration. Frontiers in Ecology and the Environment 3:161–169.

Weinstein, B. G. 2018. A computer vision for animal ecology. Journal of Animal Ecology 87:533–545.

Weinstein, B. G., L. Garner, V. R. Saccomanno, A. Steinkraus, A. Ortega, K. Brush, G. Yenni, A. E. McKellar, R. Converse, C. D. Lippitt, A. Wegmann, N. D. Holmes, A. J. Edney, T. Hart, M. J. Jessopp, R. H. Clarke, D. Marchowski, H. Senyondo, R. Dotson, E. P. White, P. Frederick, and S. K. M. Ernest. 2022. A general deep learning model for bird detection in high-resolution airborne imagery. Ecological Applications 32:e2694.

Weinstein, B. G., S. Marconi, M. Aubry-Kientz, G. Vincent, H. Senyondo, and E. P. White. 2020. DeepForest: A Python package for RGB deep learning tree crown delineation. Methods in Ecology and Evolution 11:1743–1751.

Weinstein, B. G., H. Senyondo, G. M. Yenni, E. P. White, and S. K. M. Ernest. 2024, March 17. weecology/EvergladesTools. Zenodo.

White, E. P., G. M. Yenni, S. D. Taylor, E. M. Christensen, E. K. Bledsoe, J. L. Simonis, and S. K. M. Ernest. 2019. Developing an automated iterative near-term forecasting system for an ecological study. Methods in Ecology and Evolution 10:332–344.

